# Non-polio enteroviruses in faeces of children diagnosed with acute flaccid paralysis in Nigeria

**DOI:** 10.1101/084350

**Authors:** TOC Faleye, MO Adewumi, MO Japhet, OM David, AO Oluyege, JA Adeniji, O Famurewa

## Abstract

**Background:** The need to investigate the contribution of non-polio enteroviruses to acute flaccid paralysis (AFP) cannot be over emphasized as we move towards a poliovirus free world. Hence, we aim to identify non-polio enteroviruses recovered from the faeces of children diagnosed with AFP in Nigeria.

**Methods:** Ninety-six isolates, (95 unidentified and one previously confirmed Sabin poliovirus 3) recovered on RD cell culture from the stool of children <15 years old diagnosed with AFP in 2014 were analyzed. All isolates were subjected to RNA extraction, cDNA synthesis and three different PCR reactions (one panenterovirus 5′-UTR and two VP1 amplification assays). VP1 amplicons were then sequenced isolates identified.

**Results:** 93.75% (90/96) of the isolates were detected by at least one of the three assays as an enterovirus. Precisely, 79.17% (76/96), 6.25% (6/96), 7.295% (7/96) and 6.25% (6/96) of the isolates were positive for both, positive and negative, negative and positive, as well as negative for both the 5′-UTR and VP1 assays, respectively. In this study, sixty-nine (69) of the 83 VP1 amplicons sequenced were identified as 27 different enterovirus types. The most commonly detected were CV-B3 (10 isolates) and EV-B75 (5 isolates). Specifically, one, twenty-four and two of the enterovirus types identified in this study belong to EV-A, EV-B and EV-C respectively.

**Discussion:** This study reports the circulating strains of 27 non-polio enterovirus types in Nigerian children with AFP in 2014 and Nigerian strains of CV-B2, CV-B4, E17, EV-B80, EV-B73, EV-B97, EV-B93, EV-C99 and EV-A120.

## INTRODUCTION

Enteroviruses (EVs) are members of the genus *Enterovirus* in the family Picornaviridae, order Picornavirales. There are 12 species within the genus, four (EV-A to EV-D) of which (alongside the Rhinoviruses) have been repeatedly found to infect humans. The enterovirus capsid is a non-enveloped icosahedron with a diameter of 20–30nM. It encloses a positive sense, single stranded RNA genome of ~7,500nt. The genome has untranslated regions (5′ and 3′ UTRs) flanking a single one open reading frame (ORF), whose polyprotein product is auto-catalytically cleaved into eleven proteins; four structural (VP1 – VP4) and seven non-structural (2A – 3D) proteins. Amplification of the 5′-UTR and/or the VP1 region can be used to detect the presence of enteroviruses [1–5], and the nucleotide sequence of the VP1 gene is used to identify enterovirus isolates [1–6].

Poliovirus, the best studied member of the genus *Enterovirus* and the etiologic agent of poliomyelitis belongs to species C within the genus. Humans remain the only known host of poliovirus, thus suggesting feasibility of its eradication. Consequently, in 1988, the World Health Assembly (WHA) resolved to eradicate poliomyelitis by the year 2000 [7], and the Global Polio Eradication Initiative (GPEI) was established. Courtesy of the GPEI’s activities, by year 2015 indigenous poliovirus transmission has been interrupted globally except in Pakistan and Afghanistan (www.polioeradication.org). This success has been the result of effective vaccination and intensive surveillance. Two poliovirus vaccine preparation (Oral Polio Vaccine [OPV] and Inactivated Polio Vaccine [IPV]) are currently being used by the GPEI and both are very effective [8]. However, due to the reversion of OPV, as part of the ‘end game’ strategy, the GPEI is tilting towards IPV as we approach the final phase of polio eradication [9].

Coupled with the vaccination effort is a very effective surveillance network that looks for polioviruses in both sewage-contaminated water (Environmental Surveillance [ES]) and children below the age of 15 years diagnosed with AFP. The ES strategy searches for enteroviruses in sewage-contaminated water due to the fact that all enterovirus infected individuals shed the virus in large amounts in faeces for several weeks and in turn into sewage and/or sewage contaminated water [10] (Ranta *et al*., 2001). Therefore, ES is very sensitive and can detect enterovirus isolates from both symptomatic and asymptomatic individuals. The demerit of ES based strategy is that, alone, it cannot differentiate which isolates are associated with clinical manifestations and hospitalisations. On the other hand, AFP surveillance finds enteroviruses associated with a clinical manifestations and consequent hospitalisation. However, considering that AFP surveillance detects only the <10% of enterovirus infections with clinical manifestations, the inability o f AFP surveillance to see beyond the tip of the iceberg is the strength of the ES surveillances strategy. Therefore, combining both strategies better illuminates our understanding of the epidemiological and evolutionary trajectory of enteroviruses, particularly with respect to pathogenicity. Hence, the reason the ES-AFP surveillance strategies are combined by the Global Polio Eradication initiative (GPEI) in some countries [11,12].

As part of the surveillance network are over 150 laboratories globally (Global Polio Laboratory Network [GPLN]) that use the RD-L20B isolation protocol [11,12] for poliovirus isolation. The RD-L20B isolation protocol is predicated on two different cell lines, RD (from a striated muscle cancer) (McAllister *et al*., 1969) and L20B (a murine transgenic L cell that expresses the poliovirus receptor and is consequently permissive and susceptible to polioviruses) [13, 14]. As a by-product of the poliovirus surveillance programme, the GPLN recovers several non-polio enteroviruses (NPEVs) on the RD cell line.

The earliest molecular epidemiology study documenting NPEV from AFP cases in Nigeria was carried out using the RD-L20B protocol between 2002 and 2004 [15, 16]. Subsequent studies [17–21] generating nucleotide sequence data on enterovirus diversity in Nigeria have been largely ES based. The only exception is a recent study [22] on enterovirus diversity in healthy Nigerian children, in which a cell-culture independent enterovirus detection strategy was employed. Consequently, in this study we revisit NPEVs from AFP cases in Nigeria in an attempt to revise their diversity in individuals with this clinical condition in the region.

## METHODS

### Samples

Ninety-six (96) RD cell culture isolates were analysed in this study. These isolates were recovered from the faeces of children below the age of 15 years that were diagnosed with AFP. The stool samples from which these isolates were recovered were collected in accordance with the national ethical guidelines as part of the National AFP surveillance programme in Nigeria and sent to the WHO National Polio Laboratory in Ibadan, Nigeria to ascertain whether poliovirus is the etiologic agent of the diagnosed AFP using the WHO algorithm [12]. The isolates analyzed in this study were subsequently anonymized for further studies.

The WHO algorithm stipulates that fecal suspension from all AFP cases be inoculated into RD and L20B cell lines. Isolates that show cytopathic effect (CPE) on both cell lines are considered to be polioviruses, and subsequently subjected to intratypic differentiation (ITD) to differentiate between the wild type and the vaccine strain. On the other hand, isolates that only show CPE in RD cell line are assumed to be non-polio enteroviruses and stored away since they are not of ‘urgent’ programmatic importance to the GPEI.

To assemble the 96 isolates analyzed in this study, from the archives, sixteen (16) isolates that showed CPE in RD cell line only, were randomly selected each month from January to May 2014. In June, fifteen (15) isolates were selected, and a previously identified Sabin strain poliovirus 3 was added to serve as control for the study. Besides this known isolate, none of the other 95 isolates had been previously identified.

### RNA extraction and cDNA synthesis

The algorithm followed in this study is as depicted in Figure 1. Using the RNA extraction kit (Jena Bioscience, Jena, Germany), RNA was extracted from the isolates according to the manufacturer’s instructions. Script cDNA Synthesis Kit (Jena Bioscience, Jena, Germany) was used for complementary DNA (cDNA) synthesis as instructed by the manufacturer. Specifically, random hexamers were used for cDNA synthesis as previously described [19]. The cDNA was then stored at −80°C and used for all polymerase chain reaction (PCR) assays.

**Figure 1:**
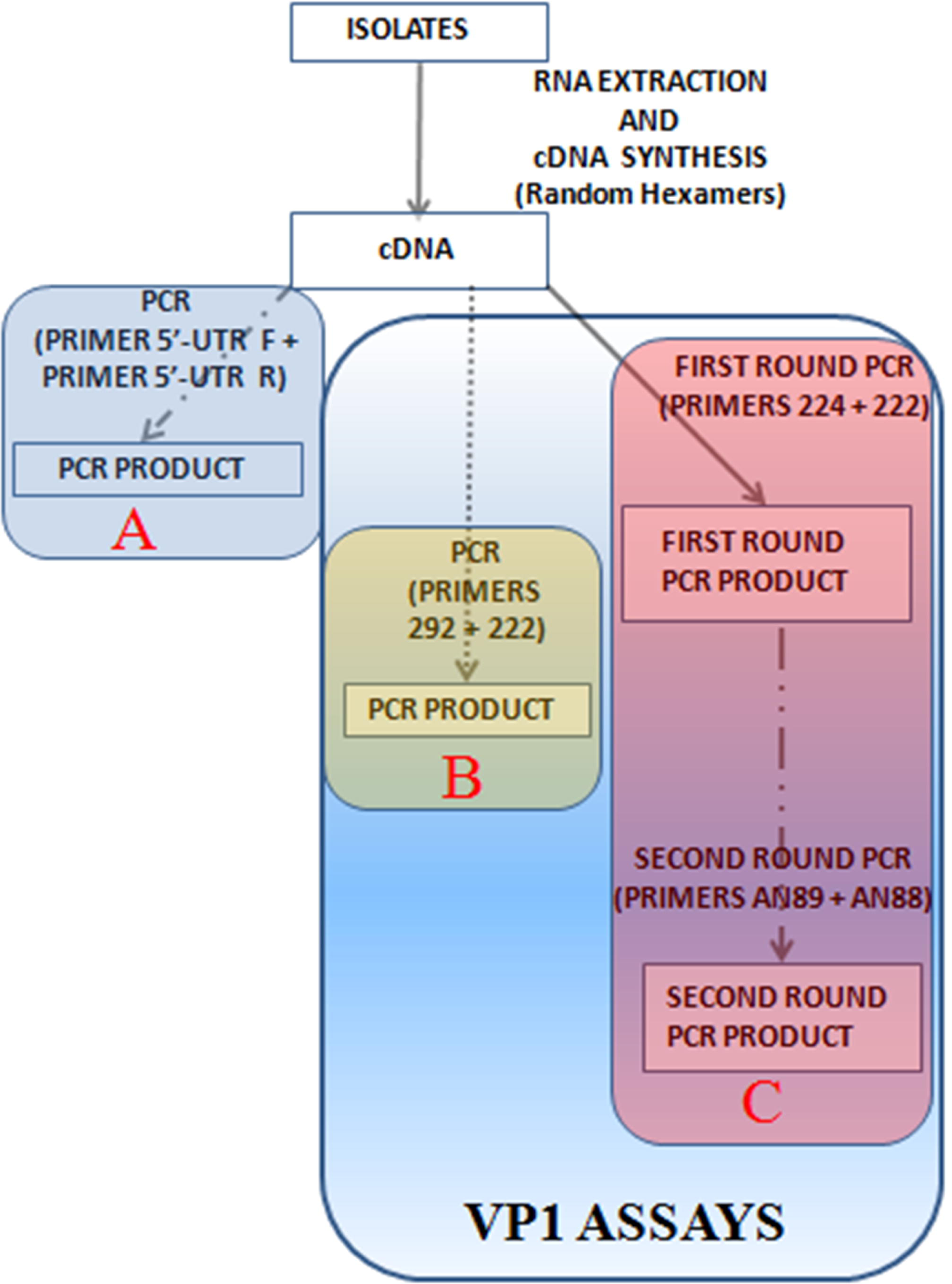
The algorithm used in this study. A depicts the 5′-UTR assay. B and C show the two VP1 assays. While B has only one stage of PCR, C has two consecutive stages (snPCR) of PCR.

## POLYMERASE CHAIN REACTION (PCR) ASSAYS

As shown in the algorithm for this study (Figure 1), three different PCR assays were run. The 5′-UTR and first PanEnterovirus VP1 PCR (PE-VP1-PCR) were one-step PCR assays. The second PanEnterovirus VP1 PCR was a semi-nested PCR assay (PE-VP1-snPCR) and was used to amplify the partial VP1 gene in those isolates for which the PE-VP1-PCR was negative. All primers were made in 100µM concentrations and all PCR assays were carried out in 30µL reaction volumes. All PCR products were resolved on 2% agarose gels stained with ethidium bromide and viewed using a UV transilluminator.

### 5′-UTR PCR and PanEnterovirus VP1 PCR (PE-VP1-PCR)

Primers panent 5′-UTR F and panent 5′-UTR R [12] were used for the 5′-UTR PCR assay while primers 229 and 222 [2, 6] were used for the PE-VP1-PCR assay. Each 30μL reaction contained 6μL of Red Load Taq (Jena Bioscience), 5μL of cDNA, 0.3μL of each primer and 18.4μL of RNase-free water. Thermal cycling was done as follows: 94°C for 3 minutes, 45 cycles of denaturation at 94°C for 30 seconds, annealing at 42°C for 30 seconds, and extension at 60°C for 30 seconds with ramp of 40% from 42°C to 60°C. This was followed by 72°C for 7 minutes, and thereafter the sample was held at 4°C until the reaction was terminated.

### PanEnterovirus VP1 semi-Nested PCR (PE-VP1-snPCR)

This assay is a semi-nested PCR assay. Primers 224 and 222 [5, 6] were used for the first round PCR while primers AN89 and AN88 [5, 6] were used for the second round PCR assay. For the first round PCR assay, the 30μL reaction contained 6μL of Red Load Taq (Jena Bioscience), 5μL of cDNA, 0.3μL of each primer and 18.4μL of RNase-free water. Thermal cycling was done as follows: 94°C for 3 minutes, 45 cycles of denaturation at 94°C for 30 seconds, annealing at 42°C for 30 seconds, and extension at 60°C for 60 seconds with ramp of 40% from 42°C to 60°C. This was followed by 72°C for 7 minutes, and thereafter the sample was held at 4°C until the reaction was terminated. The conditions were the same for the second round PCR assay except for following modifications: Instead of cDNA, the product of the first round PCR assay was used as template for the second round assay. Also the extension time for the second round assay was 30 seconds as opposed to 60 seconds for the first round assay.

### Amplicon sequencing and enterovirus typing

Only amplicon of positive PCR reactions for the two VP1 PCR assays (PE-VP1-PCR and PE-VP1-snPCR) were sequenced using the respective forward and reverse primers for each of the assays. Amplicons were shipped to Macrogen Inc, Seoul, South Korea, for purification and sequencing. Subsequently, enterovirus genotype and species were determined using the Enterovirus Genotyping Tool [23].

### Phylogenetic Analysis

The CLUSTAL W program in MEGA 5 software [24] was used with default settings to align sequences of four (the two most commonly isolated and the two for which most extensive nucleotide sequence data from the region exists) of the enterovirus serotype described in this study alongside those retrieved from GenBank. Subsequently, a neighbor-joining tree was constructed using the same MEGA 5 software with the Kimura-2 parameter model [25] and 1,000 bootstrap replicates. The accession numbers of sequences retrieved from GenBank for this analysis are indicated in the sequence names on the phylograms.

### Nucleotide sequence accession numbers

All sequences reported in this study have been deposited in GenBank and assigned accession numbers KX580638 – KX580702 and KX656918 – KX656912.

## RESULTS

### 5′-UTR assay

Precisely, 86.46% (83/96) of the isolates analyzed, including the previously identified Sabin 3, had the expected ~114bp band size and were consequently positive for the 5′-UTR PCR screen (Table 1).

**TABLE 1:** Summary of isolates characterized in this study. Twenty-seven different enterovirus types were identified in this study. Specifically, one, nty-four and two of the enterovirus types identified in this study belong to EV-A (EV-A120), EV-B and EV-C (EV-C99, SPV-3) respectively. Where icated, the asterisk (*) links the indicated amplicon to its enterovirus type, and ‘()’ denotes the number of amplicons that were successfully ntified.

| S/N | MONTH | ISOLATES<br>SCREENED | POSITIVE<br>FOR 5'-UTR<br>SCREEN | POSITIVE<br>FOR VP1<br>SCREEN | POSITIVE<br>FOR BOTH<br>5'-UTR AND<br>VP1 SCREEN | POSITIVE FOR<br>5'-UTR AND<br>NEGATIVE<br>FOR VP1<br>SCREEN | NEGATIVE<br>FOR 5'-UTR<br>AND<br>POSITIVE<br>FOR VP1<br>SCREEN | NEGATIVE<br>FOR BOTH 5'-<br>UTR AND VP1<br>SCREEN | TOTAL<br>ISOLATES<br>IDENTIFIED | SEROTYPES (Number of<br>Isolates) |
| --- | --- | --- | --- | --- | --- | --- | --- | --- | --- | --- |
| 1 | January | 16 | 15 | 14 | 13 (12) | 1 | 1 (0) | 0 | 12 | CV-B3 (2), E7 (3), E13 (2), E19 (1), E20 (1), E33 (1), E6 (2) |
| 2 | February | 16 | 14 | 15 | 13 (12) | 1 | 2 (1)* | 0 | 13 | CV-B2 (1), CV-B3 (2), CV-B4 (1), E3 (2), E12 (1), E17 (1), E20 (3)*, E21 (1), EV-C99 (1) |
| 3 | March | 16 | 14 | 13 | 13 (11) | 1 | 0 (0) | 2 | 11 | CV-B3 (1), CV-B4 (1), CV-B5 (2), E11 (2), E12 (1), E13 (1), E30 (1), EV-B75 (1), EV-B80 (1) |
| 4 | April | 16 | 13 | 14 | 12 (11) | 1 | 2 (1)* | 1 | 12 | CV-B3 (3)*, E1 (2), E11 (1), E19 (1), E21 (1), EV-B73 (1), EV-B80 (2), EV-A120 (1) |
| 5 | May | 16 | 13 | 12 | 11 (10) | 2 | 1 (0) | 2 | 10 | CV-B3 (2), E6 (2), E13 (1), E14 (1), E19 (1), E26 (1), EV-B75 (2) |
| 6 | June | 16 | 14 | 15 | 14 (11) | 0 | 1 (0) | 1 | 11 | CV-B4 (1), CV-B5 (2), E7 (1), E11 (1), E19 (1), EV-B75 (2), EV-B93 (1), EV-B97, SPV3 (1) |
|  | **TOTAL** | 96 | 83 | 83 | 76 (67) | 6 | 7 (2) | 6 | 69 |  |

### VP1 assays (PE-VP1-PC and PE-VP1-snPCR)

Fifty-nine (59) of the samples analyzed in this study, excluding the previously identified Sabin 3, had the ~350bp expected band size and were consequently positive for the PE-VP1-PCR screen. The remaining thirty-seven (37) samples were negative for the PE-VP1-PCR screen.

Of the remaining thirty-seven (37) samples, twenty-four (24), including the previously identified Sabin 3, had the ~350bp expected band size and were thereby positive for the PE-VP1-snPCR screen. The remaining 13 samples were negative for the PE-VP1-snPCR screen.

Altogether, 86.46% (83/96) of the isolates, including the previously identified Sabin 3, had the expected ~350bp band size and were consequently positive for the VP1 assays. The remaining 13.54% (13/96) were negative for the VP1 assays (Table 1).

### 5′-UTR versus VP1 assays

Overall, 93.75% (90/96) of the isolates were detected by at least one of the three assays as an enterovirus. Precisely, 79.17% (76/96) of the isolates were positive for both the 5′-UTR and VP1 assays. Furthermore, 6.25% (6/96) of the isolates were positive for the 5′-UTR assay alone, 7.30% (7/96) were positive for only VP1 assays and 6.25% (6/96) of the isolates were negative for both the 5′-UTR and VP1 assays (Table 1)

### Enterovirus Typing

Sixty-nine (69) of the 83 amplicons generated from the VP1 PCR assays and successfully sequenced were identified. The remaining 14 could not be typed due to the presence of multiple peaks in their electropherograms. Twenty-seven different enterovirus types were identified from the 69 exploitable amplicons (Table 1). Specifically, one (1), twenty-four (24) and two (2) of the enterovirus types identified in this study belong to EV-A, EV-B and EV-C respectively (Table 1).

### Phylogenetic Analysis

Of the different enterovirus types identified in this study, only four (Coxsackievirus B3 [CV-B3] and EV-B75 [the two most commonly isolated] and Echovirus 7 [E7] and E19 [the two for which most extensive nucleotide sequence data from the region exists]) were subjected to phylogenetic analysis. The CV-B3 tree (Figure 2), showed that the lineage of the CV-B3 genotype detected in 2002 (in green) is absolutely different from the lineage detected in 2014. In addition, it is important to note that the CV-B3 sequences detected in this study appear to be closely related to those from South-East Asia (Figure 2)

**Figure 2:**
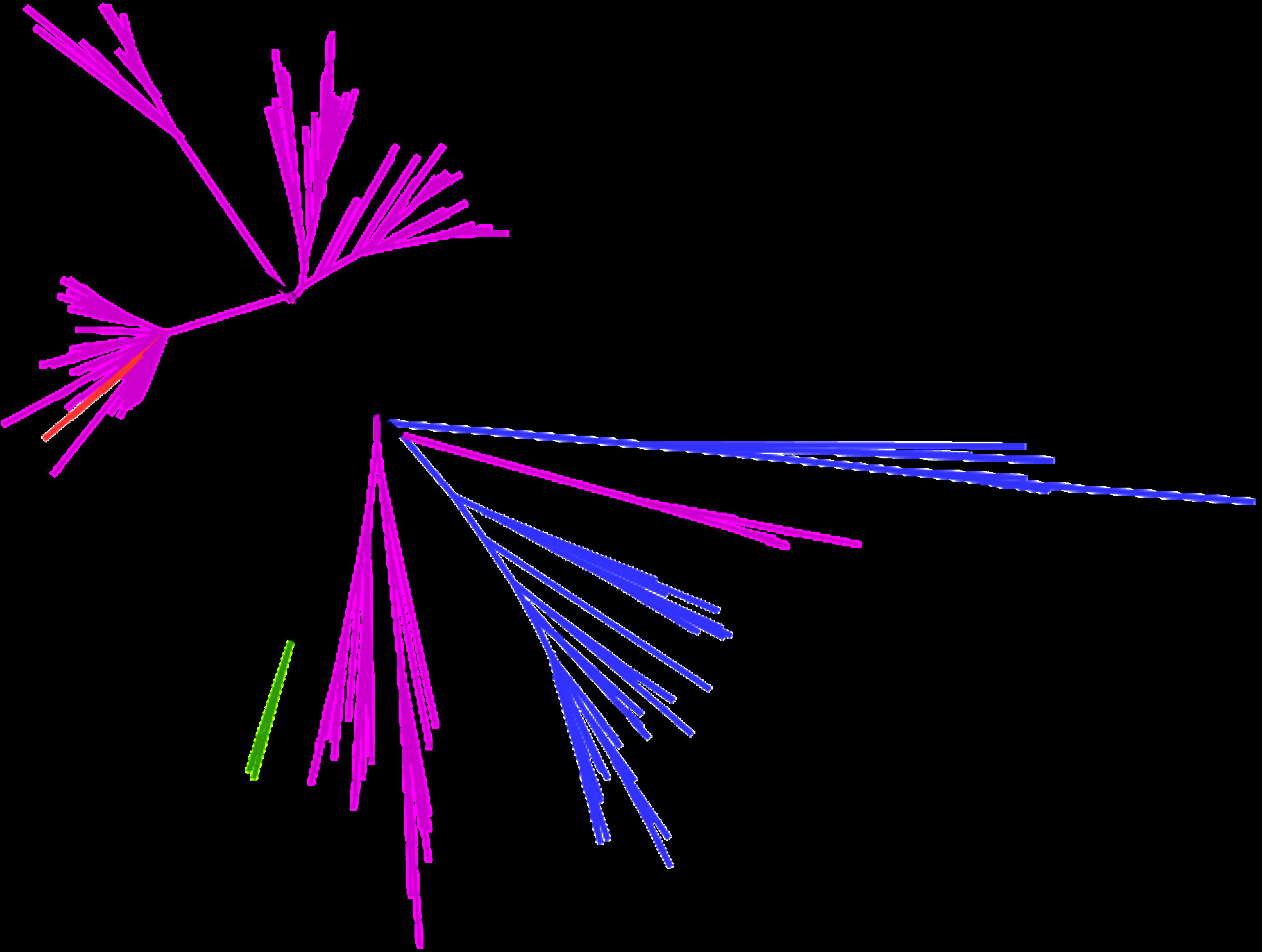
Phylogram of genetic relationship between VP1 nucleotide sequences of CV-B3 isolates. The phylogenetic tree is based on an alignment of the partial VP1 sequences. The CV-B3 sequences recovered in Nigeria in 2002 are indicated in green. The newly sequenced strains described in this study are highlighted in red. The purple clades represent CV-B3 strains for South-East Asia. The blue and black clades represent CV-B3 strains from Eurasia

For the EV-B75, two different but closely related genotypes were detected. Both were however related to EV-B75 isolates detected in Finland in 2004. One of the EV-B75 lineages found in this current study shares a common ancestor with an isolate recovered from sewage contaminated water collected in Northern Nigeria in 2012 (Figure 3).

**Figure 3:**
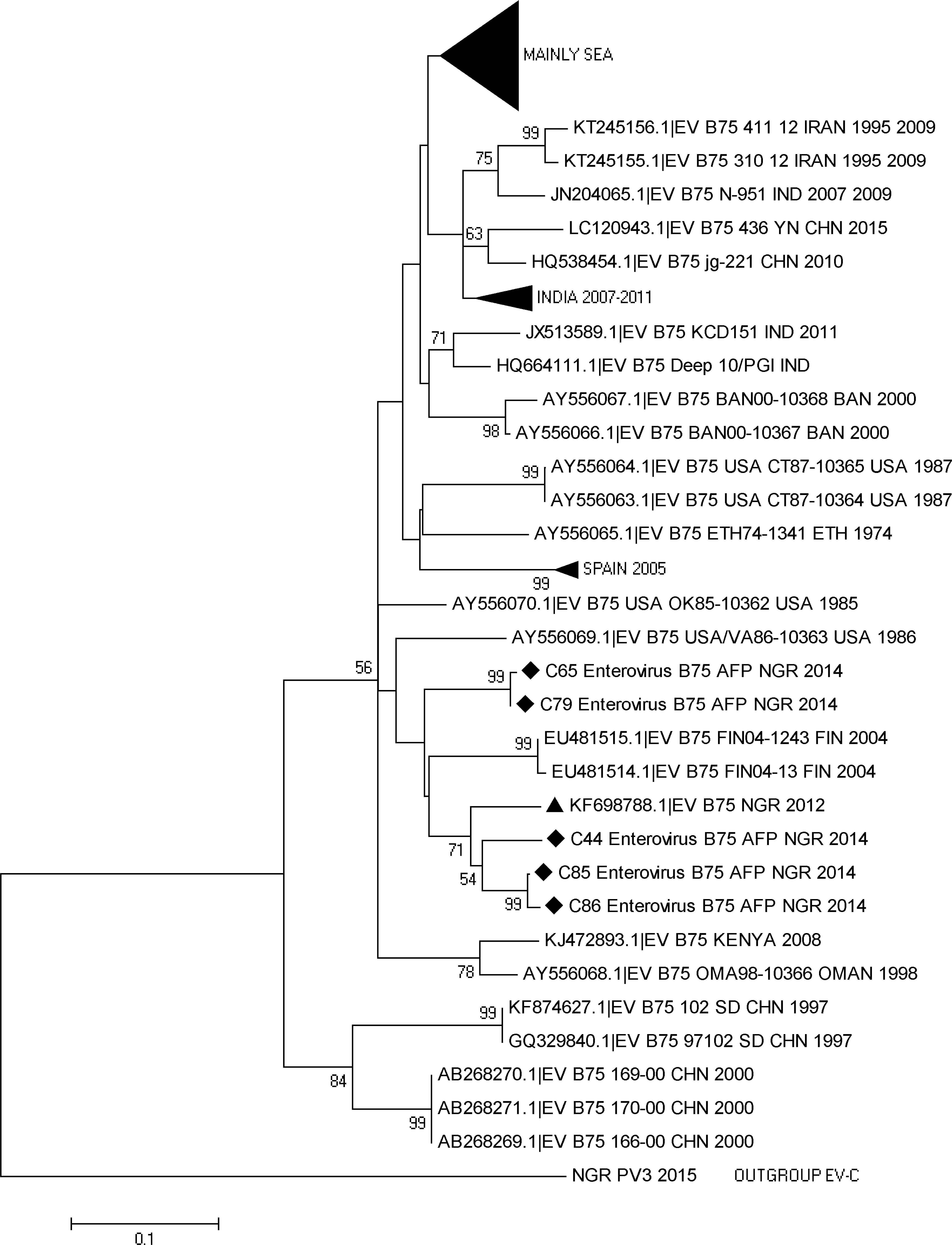
Phylogram of genetic relationship between VP1 nucleotide sequences of EV-B75 isolates. The phylogenetic tree is based on an alignment of the partial VP1 sequences. The newly sequenced strains are indicated with black diamond while the 2012 strain from Nigeria is indicated with a black triangle. The GenBank accession numbers and strain of the isolates are indicated in the tree. Bootstrap values are indicated if >50%. SEA represents South-East Asia.

For the E19, two different genotypes were detected. Both were also very different from the E19 strains isolated in 2003 (Cluster 1). The clusters to which the isolates are most closely related are indicated as 2 and 3. The single isolate from this study in cluster 2 shares a common ancestor with a lineage that was repeatedly isolated from sewage contaminated water in Lagos, Southwestern Nigeria since 2010. The remaining three strains however, belong to cluster 2 and are more closely related to an ES isolate recovered in Kano State, Northern Nigeria in 2012 (Figure 4).

**Figure 4:**
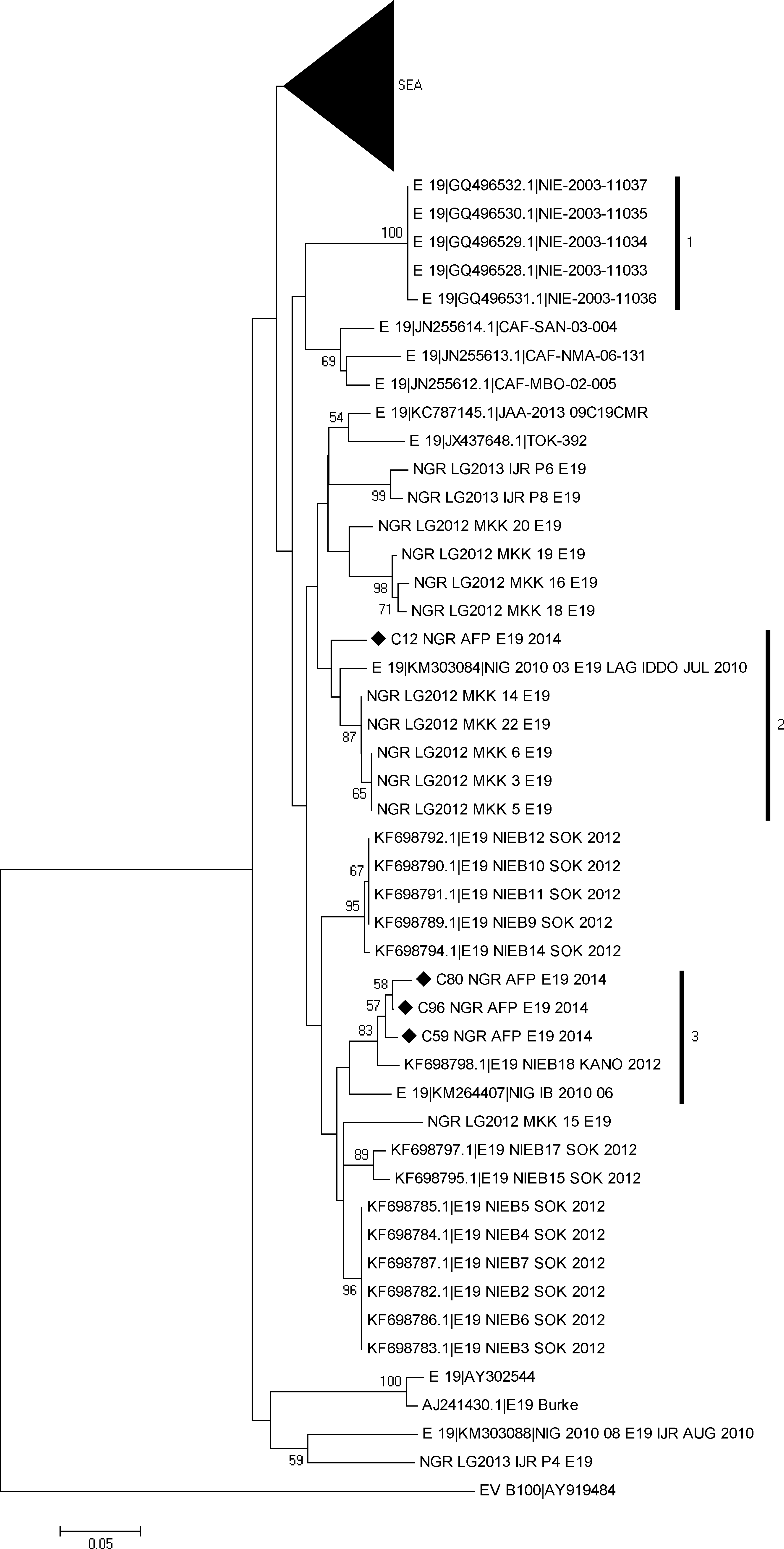
Phylogram of genetic relationship between VP1 nucleotide sequences of E19 isolates. The phylogenetic tree is based on an alignment of the partial VP1 sequences. The newly sequenced strains are indicated with black diamond. The GenBank accession numbers and strain of the isolates are indicated in the tree. Bootstrap values are indicated if >50%. SEA represents South-East Asia. The numbered vertical bars are for ease of reference alone.

As regards, E7, two different genotypes were also detected. The clusters to which the isolates are most closely related are indicated as 1 and 2. Two of the isolates from this study in cluster 1 share a common ancestor with a lineage that was first isolated from sewage contaminated water in Lagos, Southwestern Nigeria in 2010. The remaining two strains however, belong to cluster 2 and are more closely related to isolates repeatedly recovered in AFP cases in India since 2008 (Figure 5).

**Figure 5:**
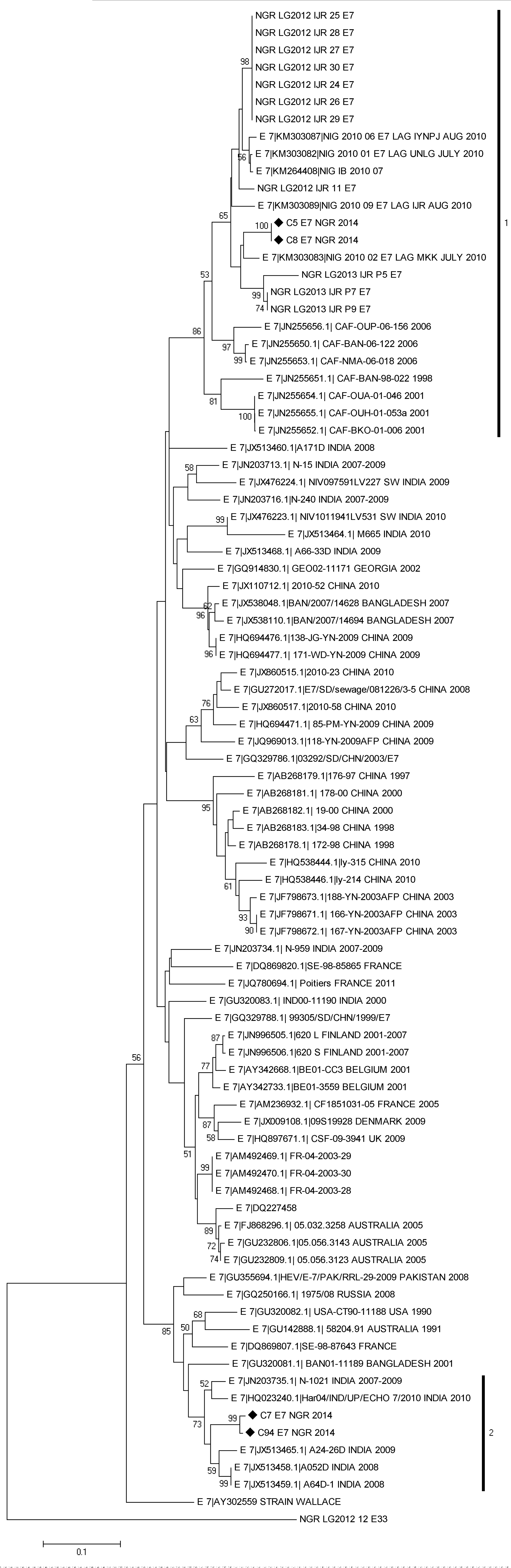
Phylogram of genetic relationship between VP1 nucleotide sequences of E7 isolates. The phylogenetic tree is based on an alignment of the partial VP1 sequences. The newly sequenced strains are indicated with black diamond. The GenBank accession numbers and strain of the isolates are indicated in the tree. Bootstrap values are indicated if >50%. The numbered vertical bars are for ease of reference alone.

## DISCUSSION

In this study, we document nucleic acid sequence data for twenty-seven (27) different enterovirus types circulating and particularly present in children below the age of 15 years diagnosed with AFP in Nigeria in 2014. It is important to mention that prior this study, no nucleic acid sequence data existed in Genbank for nine (9) of these enterovirus types from Nigeria. To be precise, to the best of the authors’ knowledge, nucleic acid sequence data for Nigerian strains of E17, CV-B2, CV-B4, EV-B97, EV-B80, EV-B73, EV-B93, EV-C99 and EV-A120 are being reported for the first time. It is however, our opinion, that the fact that these molecular sequence data are being reported for the first time does not imply new introduction of these types into the region. Rather, we believe that these types had probably been around for a long time. Hence not detecting them might reflect the lack of interest in NPEVs because the global effort focused on eradicating polioviruses. However, as the goal of poliovirus eradication nears [9], there might be an upsurge of interest in NPEVs in a bid to better understand enterovirus biology and association with varying clinical conditions.

In this study, 95.65% (66/69) and 88.9% (24/27) of the isolates and enterovirus types successfully identified, respectively, belonged to EV-B (Table 2). This is consistent with the findings of other studies from the region [15, 16, 26] predicated on the RD-L20B algorithm, irrespective of whether healthy children [26] or those diagnosed with AFP were investigated [15, 16]. However, it was recently shown that when cell-culture bias is bypassed by the direct detection of enteroviruses from stool specimen, EV-As appear to be the most preponderant in the stool of healthy children from the region [22]. Furthermore, recent studies [19, 20] showed that most CV-A13s (EV-Cs) circulating in the region selectively replicate, and can be isolated on MCF-7, but not in RD cell line. Also, when the same sample was simultaneously inoculated into RD and MCF-7 cell lines, EV-Bs and EV-Cs respectively are specifically recovered on the two cell lines [20]. In fact, we recently observed (unpublished) that when AFP samples that are negative for enteroviruses using the RD-L20B enterovirus detection algorithm [11] are subjected to the recently recommended [6] direct detection RT-snPCR algorithm described by Nix et al., [5], EV-Cs form the majority of enteroviruses detected. In addition, we also recently observed (unpublished) that about 50% of stool suspension from children with AFP in the region that yield EV-Bs in RD cell line also have an EV-C member present in the faecal suspension that was not detectable by the cell line. Put together, the preponderance of EV-B depicted by the results of this study and previous RD-L20B based studies from the region and globally should not be interpreted as an unbiased picture of the diversity landscape of enteroviruses in the AFP samples that yielded the isolates analyzed. Rather, it should be correctly viewed as the landscape as seen through the bias of RD cell culture.

Though in this study we describe enteroviruses present in the stool of children diagnosed with AFP, the findings of this study show the significance of merging ES and AFP data. Phylogenetic analysis (Figures 3, 4 and 5) suggests that representatives of some of the lineages of EV-B75, E19 and E7 detected in the AFP cases had been previously detected through environmental surveillance. Interestingly, all the sequences from Nigeria in figures 4 and 5 except those from this study and the 2002-2004 sequences were from environmental surveillance. The ES data rightly show that more lineages are circulating in the population than depicted by the AFP data reported in this study. This thereby emphasizes the power of ES to illuminate our understanding of the diversity of enterovirus types and lineages circulating in the population. This will, consequently, further enable us to better understand the evolutionary trajectory of these enterovirus types and detect their silent circulation especially in the absence of clinical manifestations.

Surveillance of enterovirus diversity among healthy children can also increase the power of the ES-AFP surveillance strategy. For instance, it was previously shown [22] that other enterovirus types not detected in this study (e.g. EV-A71 and several CV-A types) were also present and circulating in Nigeria in the same year, 2014. In addition, we had shown [22] that the EV-B80 detected in a healthy Nigerian child belonged to the same lineage as those detected in children diagnosed with AFP in this study. We further showed [22] that the EV-C99 lineage found in a healthy Nigerian child is different from that described in this study. Thus, though not suggested or included in the GPEI surveillance algorithm, enterovirus surveillance in healthy children is useful and can significantly complement the ES-AFP strategy of the GPEI.

The baseline nucleotide sequence data for CV-B3 and E19 circulating in the region were generated in 2002 and 2003 respectively [15, 16]. Hence, the data presented in this study is a re-sampling of circulating strains of these types after over a 10-year period. For both types, the strains circulating when the baseline data was generated appear to have been completely replaced (Figures 2 and 4). On the other hand, E7 and EV-B75 for which baseline sequence data from the region was generated in 2010 [17] and 2012 [21] respectively, members of the baseline lineage were still detected, however, alongside new lineages.

In some instances, it appears the variation in the population is seeded from another population. For example, as shown for CV-B3 and E7 (Figures 2 and 5), it appears that in both instances, strains from South-East Asia were imported into Nigeria and subsequently detected in the faeces of children diagnosed with AFP. These suspected importations corroborate what is known about poliovirus global circulation [8, 27]. What is not clear is why most non-polio enterovirus lineages detected in sub-Saharan Africa are yet to be detected and described in other world regions. This observation forms the basis of the regional confinement hypothesis [28].

Six (6/96) of the isolates analyzed in this study were positive for the 5′-UTR screen but negative for the VP1 screen. These isolates might actually be enteroviruses that have VP1 primer binding sites that are too divergent to be bound by the primers. Another seven (7/96) isolates on the other hand were negative for the 5′-UTR screen but positive for the VP1 assays. Should the 5′-UTR result be the basis for selecting isolates subjected to the VP1 screens, all these isolates would have been missed. It is currently not clear why these isolates were negative for the 5′-UTR screen. However, it is important to mention that recently, a new enterovirus 5′-UTR region was described that has a large deletion spanning the region amplified in this assay [29–32]. Though described in EV-C isolates, it is not clear whether such constructs are present in members of other enterovirus species and whether such constructs would be functional when transferred to other species through recombination in the 5′-UTR region. Whatever be the case, the 5′-UTR primers used in this study could not detect these isolates. Consequently, the results of this study suggests that coupling the 5′-UTR and VP1 assays might provide added sensitivity, required to find some divergent types. Particularly, it suggests that being positive for the 5′-UTR assay should not be the basis for subjecting isolates to the VP1 assays.

In this study, there were different groups of ‘untypable’ isolates (Table 1). The first group were those positive for the 5′-UTR screen but negative for the VP1 screen and have been addressed above. The second group were those positive for the VP1 assay but for which the eletropherogram could not be exploited due to multiple peaks. These are likely to come from cases where the children in question were co-infected with more than one type of enterovirus. This is not unusual and we have more recently observed (unpublished) that in about 50% of cases, stool samples from children with AFP in Nigeria contain more than one enterovirus type and/or species.

The third group were those negative for both the 5′-UTR and the VP1 assays. Considering we detected isolates that were positive for the 5′-UTR screen but negative for the VP1 screen and vice-versa, it is not difficult to conceptualize the possibility that enterovirus isolates might exist that are negative for both assays. However, currently, this is only a conjecture. Since RD cell line can also support replication of other enteric viruses like the adenoviruses [33], these third group of ‘untypables’ might not necessarily be enteroviruses but could be other viruses for which the RD cell line is both susceptible and permissive.

In conclusion, we identified 27 different enterovirus types present in Nigerian children with AFP in 2014. To the best of our knowledge, we also document the first molecular detection and identification of Nigerian strains of E17, CV-B2, CV-B4, EV-B97, EV-B80, EV-B73, EV-B93, EV-C99 and EV-A120. The results of this study suggest that coupling the 5′-UTR and VP1 assays might provide the added sensitivity, required to detect more enterovirus types. Particularly, that being positive for the 5′-UTR assay should not be the basis for subjecting isolates to the VP1 assays. The results of this study also suggest that there might be importation into Nigeria of enterovirus clades from other world regions, and especially South-East Asia. Finally, the results of this study show that surveillance of the environment, AFP and healthy children all contribute significantly towards obtaining a complete picture of the diversity of enterovirus types present and circulating in any population.

## ACKNOWLEDGEMENTS

We thank the WHO National Polio Laboratory in Ibadan, Nigeria for providing the anonymous isolates analyzed in this study. This study was funded by a Tertiary Education Trust Fund, Research Project Intervention, Abuja, Nigeria to FO. The funding body had no role in study design; in the collection, analysis and interpretation of data; in the writing of the report; and in the decision to submit the article for publication. Hence, the opinions expressed in this article are solely those of the authors.

## AUTHOR CONTRIBUTIONS

Study Design (All authors)

Sample Collection and laboratory analysis (FTOC, AMO, AJA)

Data analysis (All Authors)

Wrote the first draft of the Manuscript (FTOC)

Revised the Manuscript (All Authors)

Read and Approved the Final Draft (All Authors)

AJA and FO both supervised this study

